# Clarifying the relationship between body size and extinction risk in amphibians by complete mapping of model space

**DOI:** 10.1101/2020.10.20.347716

**Authors:** Marcel Cardillo

## Abstract

In vertebrates, large body size is often a key diagnostic feature of species threatened with extinction. However, in amphibians the link between body size and extinction risk is highly uncertain, with previous studies suggesting positive, negative, u-shaped, or no relationship. Part of the reason for this uncertainty is “researcher degrees of freedom”: the subjectivity and selectivity in choices associated with model specification. Here I clarify the size – threat association in amphibians using Specification Curve Analysis, an analytical approach from the social sciences that attempts to minimize this problem by complete mapping of model space. I find strong support for prevailing negative associations between body size and threat status, the opposite of patterns typical in other vertebrates. This pattern is largely explained by smaller species having smaller geographic ranges, but smaller amphibian species also appear to lack some of the life-history advantages (e.g. higher reproductive output) that are often assumed to “protect” small species in other taxa. These results highlight the need for a renewed conservation focus on the smallest species of the world’s most threatened class of vertebrates, as aquatic habitats become increasingly degraded by human activity.

## Introduction

Up to 54% of amphibian species may be threatened with extinction [1, 2], and current extinction rates are several orders of magnitude higher than estimated rates from the fossil record [3]. While the anthropogenic threatening processes responsible for this extinction crisis are fairly well known [2, 4–7], we are only beginning to understand how species biological traits cause different species to respond in different ways to the same threats. Identifying diagnostic biological indicators of sensitive species may allow us to recognize species not yet threatened but with a high chance of becoming threatened in the future [8], or to estimate current extinction risk in unevaluated or data-deficient species [9, 10].

Across vertebrates, one of the most consistently well-supported predictors of high extinction risk is large body size [11–14]. In amphibians, however, the relationship between body size and extinction risk is highly uncertain, with previous studies supporting positive [4, 7, 15], negative [16], u-shaped [14], or no relationship [17–19], between size and threat status. Theoretically, there are reasons to predict both positive and negative size-risk associations in amphibians. For example, larger amphibian species have longer times to maturity and hence lower population growth rates, and are more likely to be exploited for food [15, 20, 21], but smaller species have narrower geographic distributions and are more threatened by the global pet trade [21].

The varied and conflicting conclusions about the size-risk association in amphibians stems from the fact that most studies tend to analyse taxonomically and geographically restricted subsets of species, many studies rely on *p* values (which are highly sensitive to sample size) to assess the effect of body size, and different studies employ different statistical methods, and account for effects of different covariates in their models. Every published test is necessarily preceded by numerous decisions about the subset of data to analyse and the details of how the analysis should be done. While every researcher uses their expertise and judgement to make what they believe are the best possible decisions, there is a great deal of subjectivity in this process, and different researchers will have different opinions on the ideal way to set up a test of the same hypothesis.

In the psychology literature, the subjectivity and flexibility in making research decisions has been recognized explicitly, and is referred to as “researcher degrees of freedom”[22]. The premium attached to novelty in getting research published means that each published test of a given hypothesis will represent a unique combination of analytical choices. Each combination can be thought of as a sample from a very large number of possible (and theoretically defensible) combinations, the great majority of which go untested and unreported. The selective nature of published tests makes it difficult to judge the true degree of general support for the hypothesis, or (if the hypothesis does not apply universally) to understand the variation in the conditions under which the hypothesis is supported [23, 24].

An approach known as Specification Curve Analysis (SCA) aims to minimize researcher degrees of freedom by fitting all (or the great majority of) plausible, non-redundant models, and comparing the resulting distribution of effect sizes to a set of corresponding ones generated by a null model [23, 25]. The interpretation then focuses not on any particular model, but on the effect size distribution as a whole (the specification curve). This amounts to a complete mapping of model space, contrasting with the more conventional approach in biology of heuristically searching a pre-defined, highly restricted area of model space to find a single best model, or a small set of best models.

Here I use SCA, together with global databases of amphibian geography, phylogeny and biology, to clarify the association between body size and threat status in the world’s amphibians. This approach allows me to (1) assess the degree of support for any general effect of body size on extinction risk, (2) characterize the variability in the association, and (3) understand the conditions under which different sizes and directions of effect are found.

## Methods

### Sources of data

The basic model is a phylogenetic generalized least-squares regression (PGLS), which accounts for the phylogenetic covariance structure in the data. To calculate phylogenetic covariances I used the Bayesian posterior distribution of phylogenies from a recent analysis [26] which includes nearly all of the 7238 species recognized in the classification scheme of AmphibiaWeb [27]. Species-level summary values of biological variables (body size, larval development, habitat) were from the AmphiBIO database [28]. I used maximum adult body length in mm as the measure of body size, rather than body mass, because data completeness was higher (5227 species-level values for body length compared to 591 for body mass).

Species geographic distribution polygons were from the IUCN Red List [1], and these were used to derive species range sizes and centroid latitude, and to assign species to continents. Climate (mean precipitation in driest quarter) was from Worldclim [29], and the aridity variable was from the global annual average Aridity Index [30]. As a measure of human impact I used the Human Footprint, a composite index that combines human population density with measures of landscape modification, including land use and density of transport networks [31]. All spatial data were converted to raster layers of ~500km^2^ resolution and summarized as median values within the distribution of each species.

### Threat status data and treatment of Data Deficient species

The extinction risk response variable in the comparative models was an ordinal-coded transformation of the threat status classification from the IUCN Red List. Of the 7238 species in the phylogeny, 2179 are data deficient (DD). In many previous comparative analyses, DD species have been omitted from analyses or assumed to be non-threatened, but recent evidence suggests that DD amphibians may in fact be more threatened than data-sufficient species [32]. I included in the model specifications four alternative treatments of DD species: (1) omit DD species from analyses; (2) assume all DD species are nonthreatened (coded as Least Concern); (3) assume all DD species are threatened (coded as Endangered, the mid-range threat level); (4) assume the proportion of DD species that are threatened is the same as data-sufficient species (41.5%), with species assigned randomly as either Least Concern or Endangered.

### Comparative models

PGLS models were fitted using the gls function in R, with the correlation structure derived from the phylogenetic covariance matrix and phylogenetic signal in model residuals estimated by maximum likelihood. All models were run using one randomly-chosen tree from the set of 10,000 trees in the posterior distribution. Sensitivity tests showed there was very little variance in slopes and effect sizes attributable to random sampling of phylogenies from the posterior distribution (Figure 1).

**Figure 1.**
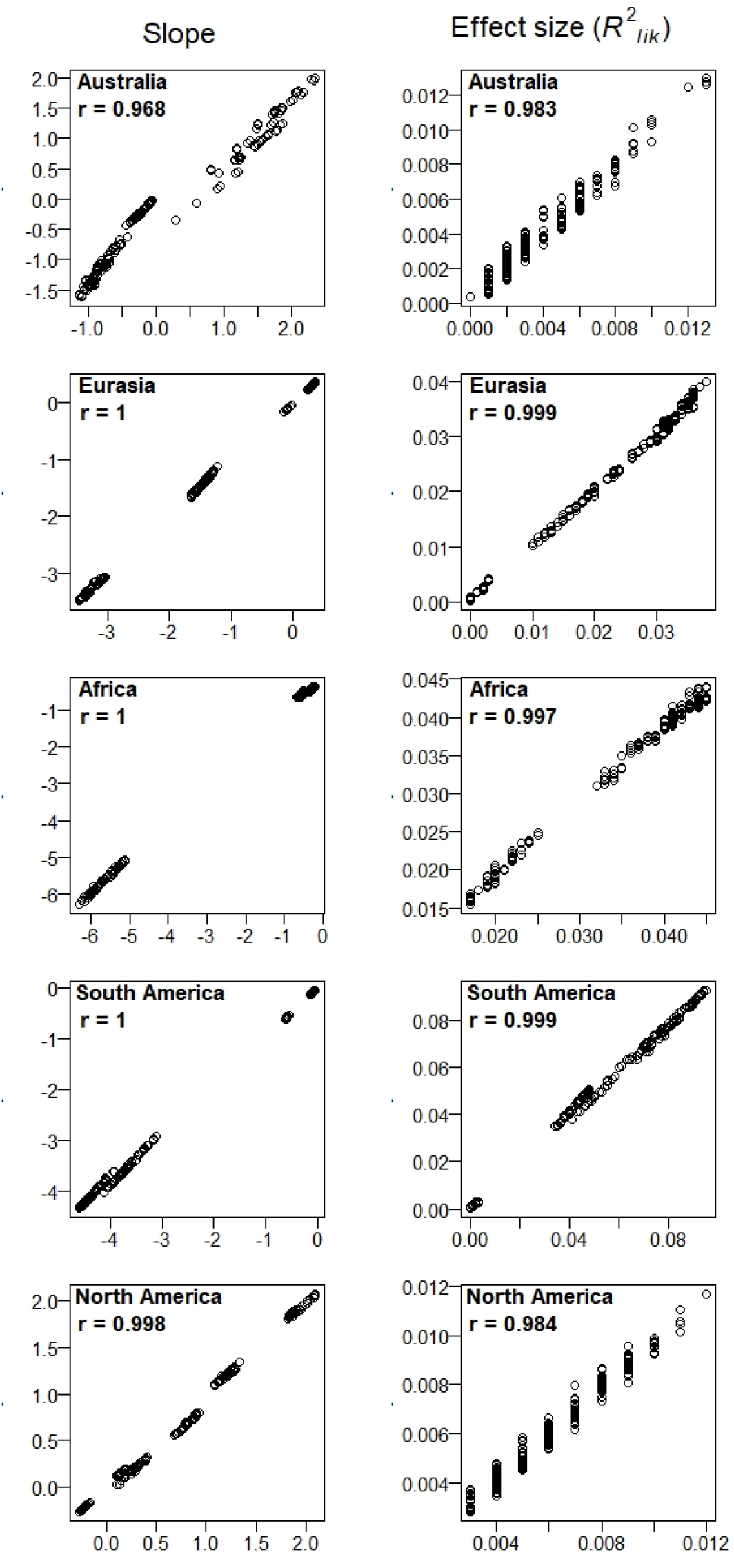
Variance in slopes and effect sizes attributable to sampling from the phylogenetic posterior set. On the x-axes of the plots are values of slopes (left column) and effect sizes (right column) for all 1024 PGLS models run on one subsampled dataset. On the y-axes are means of 50 replicate models run on the same subsampled dataset using 50 randomly selected phylogenies from the posterior set. Below the dataset name in each panel is the Pearson correlation coefficient.

With the focal variable (body length) included as a predictor in every model, there were 1024 unique combinations of the eight covariates and four treatments of DD species: this was the set of model specifications. Table 1 provides justification for considering each covariate. I divided the dataset by continent and ran models separately for each continent, to characterize the geographic variation in the body size – extinction risk association. Because of the large number of models to be fitted, and because the time taken to optimize phylogenetic branch-length transformations increases rapidly with the number of species in a model, I used a data subsetting procedure to reduce run times (which also served to hold sample sizes constant across models for different continents). From each continental dataset I chose 50 random subsets of 120 species (the number of data-complete species in North America, the minimum of any continent), and ran all 1024 models on each of these 50 subsets. The choice of 50 subsets gave estimates of mean effect sizes that were acceptably unbiased with relatively low variance (Figure 2). For the North America dataset, subsetting was not used, and only one of each model was run using the full dataset of 120 species.

**Table 1.** Justification for inclusion of covariate predictors in models, in terms of their suggested influence on the body size – extinction risk association.

| Covariate | Justification | References |
| --- | --- | --- |
| Body length<sup>2</sup> | Recent evidence for nonlinear body size – extinction risk association | 14, 20 |
| Larval developmental stage | Aquatic larvae may be more susceptible to Bd and other waterborne pathogens, and more exposed to water pollution | 38 |
| Aquatic foraging habitat | Freshwater species less protected by “faster” life history; Smaller species more likely to have fully aquatic life cycle | 27 |
| Geographic range size | Strong predictor of threat status in its own right; smaller species tend to have smaller ranges | 4, 16 |
| Latitude | Bd infection more prevalent in warm, wet climates; possible size-specific susceptibility to infection | 6, 22, 23 |
| Mean precipitation driest quarter | Larger size may offer defence against desiccation during seasonal drought | 18 |
| Aridity Index | Larger size may be desiccation defence in hot & dry climates | 18 |
| Human Footprint Index | Heavily modified landscapes may favour the lower dispersal capabilities of smaller species; More small species occupy aquatic habitats, which have been heavily modified | 5, 27 |

**Figure 2.**
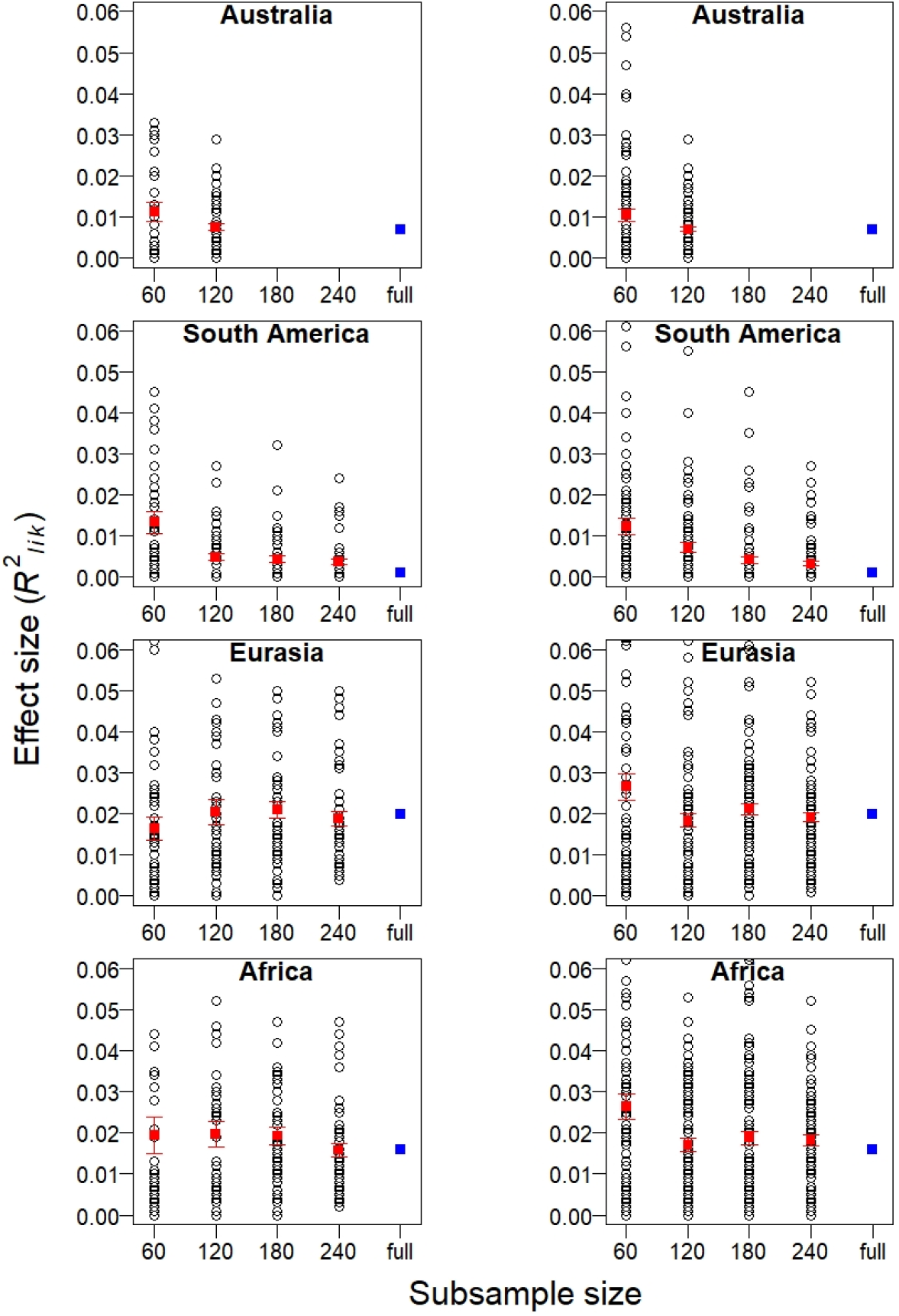
Effect of random subsampling on estimates of body size effect size against threat status. Effect sizes are partial *R*^2^_lik_ values from a PGLS model with log(body length) as the sole predictor and data deficient species omitted. Effect sizes are shown for the full dataset (blue squares; Australia: n = 145; South America: n = 1407; Eurasia: n = 491; Africa: n = 504), and subsamples of 60, 120, 180, and 240 species, and for sets of 50 (left column) and 100 (right column) subsamples. The red points and bars are the mean and standard errors of effect sizes within each set of samples.

For each model, the effect size of body size against threat status was calculated using Ives’ *R*^2^_lik_, a partial *R*^2^ statistic devised explicitly for use with models that include a covariance structure [33]. This statistic is formulated as a comparison between a “full” and “reduced” model, so to calculate the partial effect of body size I compared each model with a corresponding model in which values of body size were randomized, but the response and all other predictors left intact.

### Specification curves and null models

For each of the 1024 model specifications, I obtained the mean and standard error of *R*^2^_lik_ for the 50 random data subsets (with the exception of the North American dataset, for which only the full dataset was used). For models in which slope of the body size main effect was negative, I assigned a negative sign to the value of mean *R*^2^_lik_, in order to distinguish negative and positive effects for visualization purposes. I then plotted the effect size means in rank order, from smallest to largest, to construct the specification curve. To generate a null model for comparison, I re-fitted each model 500 times with the response variable shuffled, and recorded whether the body-size effect of each model fell within the lower or upper tails (0.025 and 97.5 quantiles) of the null distribution of 500 effect sizes for that model.

## Results

Specification curves show the distribution of ranked effect sizes of body size on threat status, for each continent (Figure 3). For the three largest datasets (Africa, Eurasia, South America), 97-99% of models indicate a negative association between body size and threat status. Although effect sizes are relatively small in all models (partial *R*^2^ < 0.05), many of the models have body size effects larger than expected under null models (indicated by red bars in Figure 3). In the smaller datasets (Australia, North America), on the other hand, body size seems to have little association with threat status: effect sizes are all <0.02 and none of these models have effects significantly greater than null expectations.

**Figure 3.**
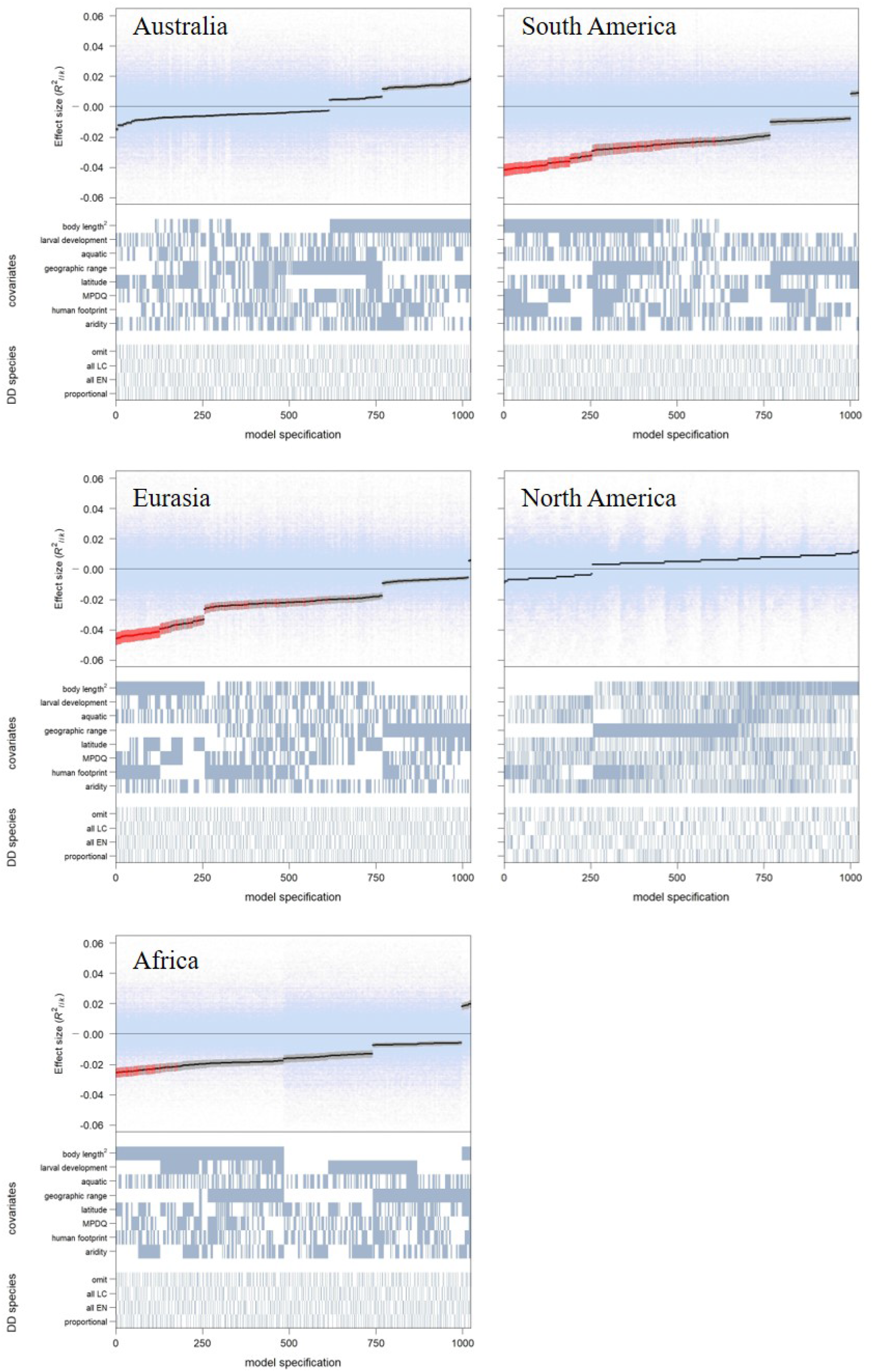
Specification curves for 1024 models of body size (log adult body length) against IUCN threat status for the world’s amphibian species, partitioned by continent. For each model, the upper panels indicate the mean (black dots) and standard error (grey bars) of the partial effect of body size for a set of 50 data subsets (n=120). Light blue points are effect sizes for 500 null-model replicates for each model specification. Means and error bars are shown in red where the mean ± standard error lies in the upper (>0.975 quantile) or lower (<0.025 quantile) tail of the corresponding null distribution. Lower panels show the set of covariates included and the treatment of data deficient species for each model specification.

The panels below each curve in Figure 3 show the specification of each model in the curve, and reveal the analytical choices that influence the size and direction of the effect of body size on threat status. In the South American, Eurasian and African datasets, the strongest negative effects are seen when a quadratic term is included in the models, so that explanatory power is greatest when the association between body size and threat status is assumed to be non-linear. When the main effect slope is negative the quadratic term is nearly always positive, implying either a parabolic association with lowest threat status for mid-sized species [14], or a negative asymptotic association with little body size effect among species of larger size. In the same three continents, accounting for the effect of geographic range size produces less negative effects of body size. In South America and Africa, including a measure of human impact (Human Footprint Index) leads to stronger negative body size effects on threat status. Other covariates, and decisions about the assumed threat status of data deficient species, have little consistent influence on the body-size effect.

## Discussion

The prevailing negative association between body size and extinction risk in amphibians is at odds with the body size effect generally found in other vertebrate groups, in which the largest species tend to be the most highly threatened. This may reflect the generally small size of amphibians compared to other vertebrates, their atypical ecology, and the particular anthropogenic impacts that threaten them. The specification curves show that the body size effect on threat status tends to be less strongly negative in models that include geographic range size as a covariate (Figure 3). Previous global-scale studies have also found that the size – threat association in amphibians is mediated by range size in a similar way [4, 16] (although both of these studies found that the association switches from negative to positive when range size is accounted for). This would suggest that range size captures much of the negative association between body size and threat status. This pattern can be explained by strong positive associations between body size and range size (Table 2), implying that small species are frequently limited to narrow distributions, leaving them more vulnerable to habitat loss and other disturbances.

**Table 2.**
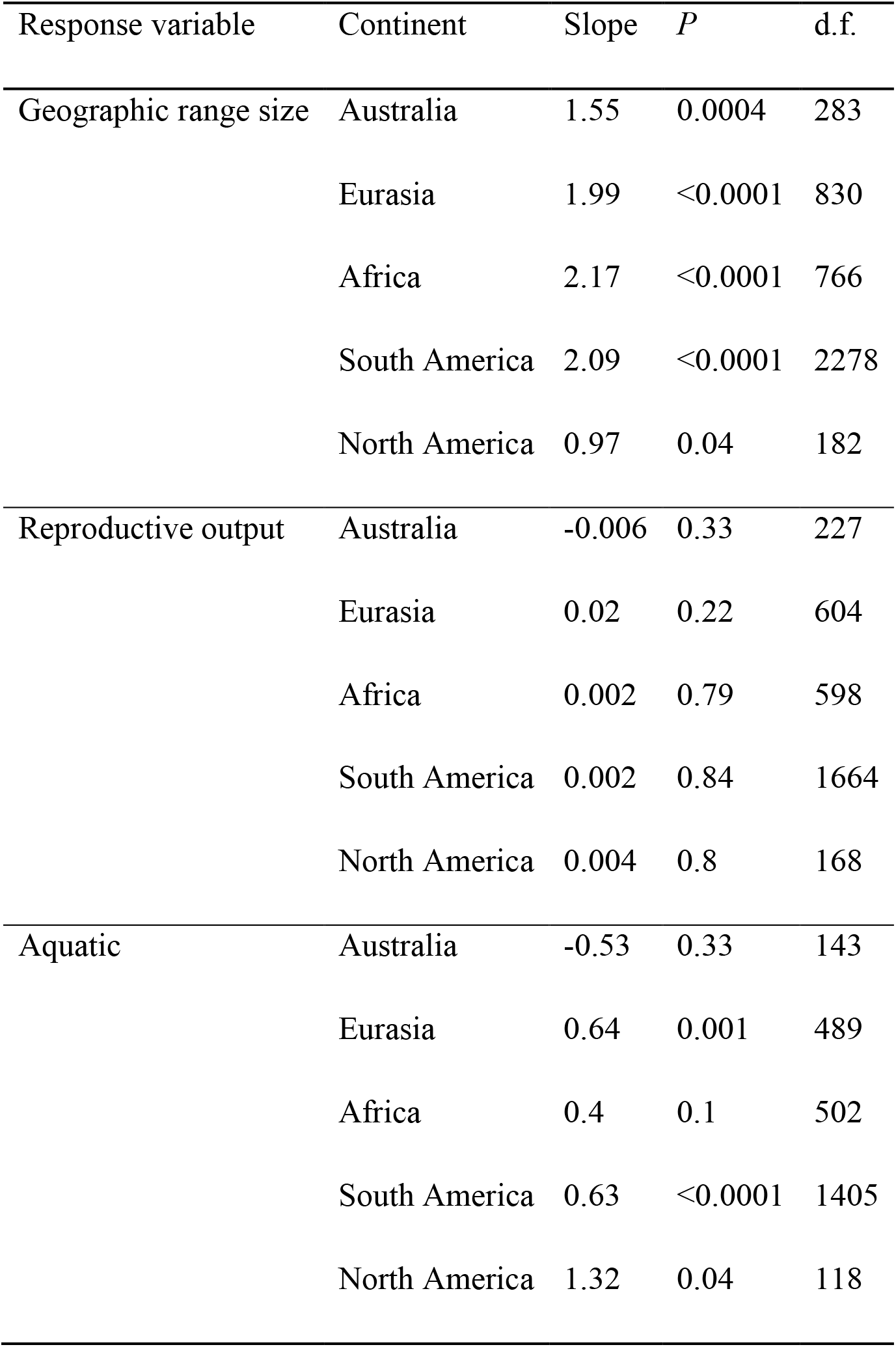
Associations between body size and geographic range size, reproductive output (maximum number of reproductive events/year), and aquatic habitat, for amphibian species in each of five continents. For all models, the predictor variable is log(body length). Geographic range size and reproductive output are log-transformed. Models are PGLS for range size and reproductive output, phylogenetic logistic regression for aquatic habitat.

However, a positive association between body size and geographic range size also holds true for vertebrate taxa other than amphibians, in which we typically see positive size – threat associations. If range size strongly mediates the effects of body size on threat status, then we need to explain why we still see mostly negative size – threat associations in amphibians, even after controlling for the effects of range size. This pattern suggests that other features associated with small size in amphibians also elevate extinction risk, independently of geographic range size.

Uniquely among vertebrates, amphibians have suffered a highly destructive global disease outbreak, with the chytridiomycosis panzootic resulting from the fungal pathogen *Batrachochytrium dendrobatidis* (Bd) responsible for population declines of at least 500 species and extinction of 90 species worldwide [6]. However, Bd is unlikely to be the primary driver of a negative body size – extinction risk association. On a global level, larger-bodied species appear to have been more severely affected than smaller ones [6], possibly because the larger size and longer lifespan of larger individuals allows them to develop a higher pathogen load [34, 35], although increased sensitivity of smaller individuals to Bd has also been reported [34, 35]. Furthermore, the effects of Bd are not yet global, and are still limited in some continents, including Eurasia and Africa [6], where negative size – threat associations also prevail. It therefore seems likely that the higher threat status of small-bodied species is the result not of chytridiomycosis, but of threatening processes that are even more pervasive and widespread globally.

Another possible explanation is that amphibian species’ strong dependence on freshwater ecosystems, with most species having an aquatic stage for at least part of their life cycle, represents an additional ecological disadvantage for amphibians [36]. Globally, freshwater ecosystems have been disproportionately affected by anthropogenic disturbances, including invasive species, pollution, wetland drainage and river flow modification [37, 38], imposing additional pressure on populations of amphibians, in a way that may compound the negative effects of a small distribution. Again, however, this explanation does not seem likely to account for prevailing negative size – threat status associations in amphibians, for several reasons. In the data I have used in this study, the disadvantage of being dependent on freshwater ecosystems is best captured by two variables: aquatic foraging habitat (a variable from the Amphibio database) and human impact, represented by the Human Footprint Index. If a dependence on freshwaters was partly responsible for the disadvantage of being small, we would expect to see a reduction in the magnitude of the body-size effect in models where either or both of these variables appear as covariates, but this is not what we see in the specification curves (Figure 3). Furthermore, the global amphibian dataset suggests that it is in fact larger, not smaller, species that tend to be more strongly associated with aquatic ecosystems. Phylogenetic logistic regression models indicate strong positive associations between aquatic habitat and body size for most continents (Table 2).

Another possible contributing factor to the prevailing negative size – threat associations is that for amphibians (unlike many other vertebrate taxa) there is simply little advantage in smaller size, in terms of the speed of life history and intrinsic rate of population growth. A recent study has shown that body size is less important in determining life-history speed in amphibians compared to reptiles, probably because the range of body sizes represented by amphibian species is more limited [39], and this is likely also to be true in comparison to other vertebrate taxa. The Amphibio database supports this conclusion: there are no significant associations between body size and reproductive output, the life history variable with the greatest coverage in the database (Table 2).

In conclusion, it seems that smaller amphibian species tend to have higher extinction risk than larger species, largely by virtue of their smaller geographic distributions, but it remains unclear whether additional factors also predispose smaller species to higher risk independently of range size. Although effect sizes are small, the association between smaller size and elevated risk in amphibians is clear and globally widespread, and robust to a wide variety of model formulations. Given the evidence in many vertebrate taxa that high extinction risk is associated with large size, and the high conservation profile of many large-bodied threatened species, it is often assumed that smaller species – accounting for the bulk of animal diversity – are afforded protection by their faster life history (e.g. larger litters or shorter times to maturity) and high rates of population increase. The results of this study call into question the generality of this assumption, and suggest that it may be a dangerous assumption for amphibians. In this context, the results of this study lend support to recent calls for a renewed focus on the conservation needs of the smallest amphibians [36].

To the best of my knowledge, this study is the first application of the SCA approach to exploring and testing a hypothesis in conservation science. There are two key strengths of this approach of complete model-space mapping. The first is transparency – by presenting the results of all plausible models, we avoid the pitfalls of researcher degrees of freedom, in which subjectivity and opinion play a strong role in defining the bounds of model space to be explored, and in the selection of models to interpret from within this space [22]. Nonetheless, even the process of defining the “universe” of plausible models has a certain degree of subjectivity: for example, the set of covariates to include were those I considered to have *a priori* justification for examining the effect on threat status of amphibians, but other researchers may have chosen to exclude some of these or include others. The second key strength of the SCA approach is that it reveals the full range of variation in the form and strength of the pattern being tested, and allows us to discover the conditions under which the hypothesis is supported, or is not supported.

## Acknowledgements

I am grateful to Ben Scheele, Renee Catullo, Craig Moritz, Octavio Jiménez Robles, Ian Brennan, and members of the Macroevolution & Macroecology Group, Australian National University, for helpful comments and feedback. This research was supported by the Research School of Biology, Australian National University.

